# Evaluating a SNP calling pipeline for *Mycobacterium leprae*

**DOI:** 10.1101/2025.07.18.665481

**Authors:** Katherine Cox, Daniel Whiley, Aureliana F.C. Chilengue, Maria Rosa Domingo Sananes, Conor Meehan

## Abstract

*Mycobacterium leprae* is the causative agent of leprosy, a persisting disease characterised by skin lesions and peripheral nerve numbness. Attempts to sequence the genome have been challenged by its inability to be cultured on artificial media. Advances in DNA extraction and sequencing technologies have enabled molecular epidemiology to be introduced for *M. leprae*. However, a validated SNP calling pipeline does not currently exist. A ground truth of pairwise SNP differences was first created using Minimap2 and three complete genomes of *M*. *leprae*. Using simulated reads from each of these genomes and the others as a reference, we evaluated the precision and recall of short-read SNP calling using Snippy. Approximately 80% of SNPs were called with false positives only due to ambiguous bases in certain genomes. Repeat region masking was found to be unnecessary for *M*. leprae SNP calling, unlike for *M*. tuberculosis. We find that SNP calling from short reads is robust and highly accurate from *M. leprae*, showing promise as a tool for molecular epidemiology studies to increase case detection and inform leprosy control strategies.

## 1. Introduction

Leprosy is a chronic infection of the skin, peripheral nerves, eyes and respiratory tract caused primarily by *Mycobacterium leprae* [1]. It is characterised by hypopigmented, insensitive skin lesions and numbness in the extremities. Secondary infections, injuries and progressive nerve damage can lead to tissue loss, deformities, and paralysis of the hands and feet [1,2]. The average incubation period for leprosy is around 5 years but can extend over several decades [4-6]. Leprosy persists as a neglected tropical disease (NTD), predominantly affecting low-to middle-income countries (LMICs) [4]. Approximately 200,000 new cases are reported each year [4] and are largely concentrated in India, which reports over 100,000 cases annually [7], as well as Brazil and Indonesia, each reporting over 10,000 [8].

While the transmission of leprosy has been linked to aerosols and prolonged close contact with infected individuals, the underlying mechanisms remain unknown [9]. Eradication relies on identifying transmission clusters to inform resource allocation and public health interventions [10]. Following its elimination as a global health concern, there has been a decline in case detection activities [11,12]. As such, studies predict the true number of cases is much higher than those reported [11,13]. The disability and stigma associated with advanced infections discourage many from reporting their illness [14]. Additionally, healthcare may be inaccessible or inadequate in poor and remote regions [2], and the lesions caused by leprosy may be confused for other, more common skin conditions [15].

The genomic tracking of pathogens can enhance case surveillance, revealing transmission links which may be missed through clinical reporting alone [16]. However, molecular studies of *M. leprae* have faced their own challenges, with advances only seen in the last two decades. For many years, efforts to sequence *M. leprae* were hindered by the nature of its genome, up to 60% of which consists of pseudogenes as a result of reductive evolution [17]. The absence of functional metabolic pathways leaves the organism reliant on a host to provide nutrients [18]. Consequently, *M. leprae* is almost completely unculturable *in vitro*, complicating antimicrobial susceptibility testing and whole genome sequencing (WGS) efforts [6]. The discovery of the nine-banded armadillo [19,20] and, later, the mouse footpad [21,22] as viable hosts to cultivate *M. leprae* led to significant breakthroughs in the understanding of its genome [17,18].

The first *M. leprae* genome was completed in 2001 from a strain isolated in Tamil Nadu, India. This strain, known as the TN strain, was determined to be 3.3Mb long [17]. In the same study, TN was compared with *Mycobacterium tuberculosis* strain H37Rv, revealing extensive genome reduction. In the two decades since, only three more genomes have been completed [23]. Comparative analysis in 2005 between the TN strain and a 142 Kb portion of the Br4923 Brazilian strain found the *M. leprae* genome to be highly conserved, with a single nucleotide polymorphism (SNP) frequency of around 1 per 28 kilobases detected [18]. In 2009, analysis of the TN strain, the now completed Br4923 strain and two strains from Thailand and the United States revealed over 99% sequence identity [23]. In 2018, a genetic substitution rate of 7.8×10^−9^ was reported, indicating a remarkably slow evolution [24].

The 2005 comparison of the TN strain with the limited Br4923 genome revealed three key informative SNPs. When applied to a dataset of 175 samples across 21 counties, four SNP types were identified and designated 1 to 4 [18]. The subsequent comparison of TN with the complete Br4923 produced 78 informative SNPs, 6 insertions/deletions (indels) and four homopolymeric tracts. 400 isolates from across the globe were analysed for these features, resulting in the identification of 16 SNP subtypes [23]. This was later updated to 19 with the discovery of new genotypes 1D-Malagasy [25], 1B-Bangladesh [26], 3K-0, 3K-1 and 4N/O [24]. Each of these studies reported a strong correlation between SNP type and geographical location, providing insights into the global and historical dissemination of leprosy [6].

The SNP typing system only focuses on 78 informative SNPs, often widely considered too conserved for fine-scale epidemiological investigations [18,23]. Conversely, variable number tandem repeats (VNTR) were found to exhibit extreme variation – even between samples taken from the same patient [27]. VNTR is therefore recommended for smaller scale studies and SNP typing for widespread epidemiological investigations [7]. However, comparative analysis of all available strains revealed a total 4,770 SNPs [28] which could provide further insights into the *M. leprae* genome. Moving beyond the original 78 SNPs and examining SNP distances may reveal a wealth of information regarding the drug resistance, transmission and phenotypic variations which impact disease manifestation [28]. Thus, the use of SNP calling in clinical molecular epidemiology studies could be a powerful approach to enhance case detection and inform leprosy control strategies [24,25]. Currently, no SNP calling framework for *M. leprae* exists, leaving its potential untapped.

Advances in DNA extraction and sequencing technologies mean that direct-from-sample sequencing is becoming more common [7,24], predominantly from MB leprosy cases due to the moderately higher bacterial load in the skin lesions [1]. This process offers a faster alternative to the traditional animal passaging methods which take at least 6 months due to the 12-day generation time of *M. leprae* [6]. As the amount of available data increases, the need for a clinical WGS pipeline to resolve transmission clusters becomes evident. It is imperative to remember that high burden, low resource regions have the most immediate need. We therefore evaluated the accuracy of a low cost, easy to use SNP calling pipeline which could guide future molecular epidemiology studies and leprosy control measures.

## 2. Materials and Methods

### 2.1. Assessment of available complete genomes

All complete assembled M. leprae genomes were downloaded from the National Centers for Biotechnology Information (NCBI) genome database [33]. To aid the later interpretation of results, the completeness of these genomes was evaluated using BUSCO version 5.8.3 [34,35]. The mode was set to genome and auto-lineage for prokaryotes.

### 2.2. Establishment of ground truth SNPs for pipeline validation

Genome alignment was performed on each aforementioned complete genome using Minimap2 version 2.28 [36,37] with asm5 used for < 5% sequence divergence. Penalties of 20 and 60 were introduced for opening and extending gaps, respectively. The program was run pairwise, with each comparison producing a variant call format (VCF) file containing every detected SNP and its position, along with any gaps introduced. The SNPs and their positions served as the ‘ground truth’ which read mapping approaches would be compared against for validation.

### 2.3. Simulation of short read sequences and read mapping

Paired-end reads were simulated from the original complete genomes using InSilicoSeq version 2.0.1 [38] set to model Illumina MiSeq sequencing with a read depth of 100.9, an average genome length of 3.31Mb and a read length of 300 base pairs (bp). Pigz version 2.8 [39] was used to rapidly compress the output files.

Read mapping was performed using Snippy version 4.6.0 [40] with default settings. Similar to Minimap2, Snippy was run pairwise with each pair of short reads mapped to each complete genome as a reference. The VCF output contained SNPs, insertions, deletions, complex regions and multiple nucleotide polymorphisms along with the positions at which they occurred.

### 2.4. Simulation of realistic read lengths using available clinical samples

All NGS samples listed as *M. leprae* in the European Nucleotide Archive (ENA) database [41] were downloaded on 16th December 2024. Snippy was run on each genome with default settings using one of the complete genomes as reference. Pandepth version 2.25 [42] was used to filter out any samples with < 90% genome coverage.

FastQC version 0.12.1 [43] was run on the remaining samples to generate quality reports. These were collated using MultiQC version 1.27.1 [44] and analysed to determine read lengths most commonly found in clinical datasets. These parameters were used to simulate realistic read lengths with InSilicoSeq as above. Reads were again compressed using Pigz, and Snippy was run on each set as described in section 2.3.

### 2.5. Estimation of model performance

Using Python version 3.8.10, the VCF files containing the SNPs and positions identified by Snippy were compared to the VCF ground truth files generated by Minimap2. The true positive (TP), false positive (FP) and false negative (FN) SNPs were listed in text files along with their positions and proximity to other SNPs and gaps. The number of TPs, FPs and FNs were used to calculate precision, recall and F1 score metrics [31]. Precision measures the proportion of predicted SNPs which are true. It is calculated using the number of TPs and FPs in the calculation:

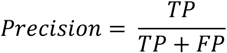

Recall is the proportion of true SNPs which have been predicted by the model and is calculated using TPs and FNs in the calculation:

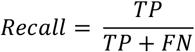

The F1 score balances precision and recall to ensure that as many SNPs as possible are correctly identified without introducing errors (FPs). This is calculated using precision and recall values in the calculation:

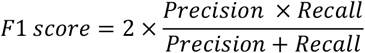

These metrics were then used to evaluate the performance of Snippy and the quality of each genome as a reference. Where FP SNPs were called, these were analysed using Mauve version 2.4.0 [45].

### 2.6. Masking of low complexity and repeated genome regions

Genmap version 1.3.0 [46] was used to mask regions in each genome which repeat within a given k-mer length. A k-mer length of k=50 was selected to improve sensitivity while minimising over-masking when applied to short reads. Bedtools version 2.31.1 [47] was used to merge any gaps smaller than the 50-base k-mer length and generate a final browser extensible data (BED) file. Low complexity regions were identified using AlcoR version 1.9 [48] with default settings, producing a text file of the location and length of masked regions.

The Snippy VCF files were masked using the resultant BED files from Genmap and AlcoR. These were again compared to the Minimap2-produced ground truth as described in section 2.5 to determine whether masking these regions affects read mapping and SNP calling.

### 2.9. Data visualisation and analysis

Plots were generated for data visualisation in GraphPad Prism 10 [51] and RStudio version 2023.06.2 [52] using R version 4.3.1 [53]. The packages used included dplyr version 1.1.4 [54], GGally version 2.2.1 [55], ggplot2 version 3.5.1 [56] and tidyr version 1.3.1 [57].

Statistical analysis was performed in GraphPad Prism 10.4.2. A confidence interval of 95% (P < 0.05) was used for all tests. One-way analysis of variance (ANOVA) tests were used to determine whether the difference in results across read lengths and approaches were statistically significant. A post-hoc Bonferroni correction was applied to account for multiple tests.

## 3. Results

### 3.1. Genome completeness

There are currently only four complete genomes available on NCBI. These are of the Indian strains TN and MRHRU-235-G, along with Japanese strain Kyoto-2 and the Brazilian strain Br4926. When analysing these genomes through BUSCO, the *Actinomycetes* lineage was detected as a best match for comparison. Of the 355 total orthologs, an average 92% were present in each genome in a single copy. 6.6% were missing, and 1.4% were fragmented (Figure 2). No orthologs were present in more than one copy.

**Figure 2.**
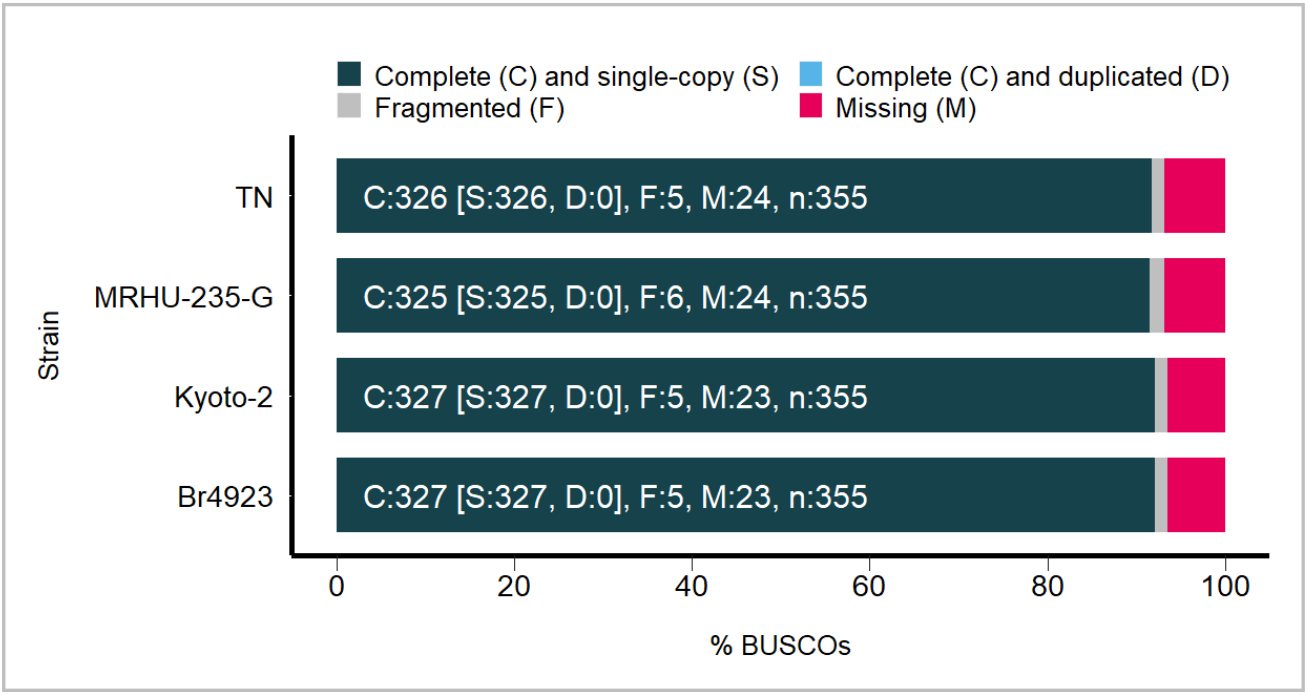
The results of BUSCO genome assessment using the *Actinomycetes* lineage. Each bar represents a complete genome and is split by colour into percentages of the total number (n) of orthologs in this lineage. Dark blue corresponds to complete (C) single-copy (S) orthologs, while light blue corresponds to complete duplicated (D) orthologs. Grey represents fragmented (F) orthologs and pink represents the proportion of orthologs missing (M). The graph was created and adapted using R script generated by BUSCO.

### 3.2. Ground truth SNPs

The SNPs between each complete assembled genome as identified by Minimap2 are shown in table 1. These numbers are the ‘ground truth’, providing the frame of reference for subsequent results. The distance between genomes ranged from a minimum of 110 SNPs and a maximum of 249.

**Table 1.** The number of SNPs identified between each aligned pair of genomes by Minimap2. These figures serve as the ground truth to compare other approaches and pipeline tools.

| Reference | Query | SNP distance |
| --- | --- | --- |
| Br4923 | TN | 155 |
| Br4923 | MRHRU-235-G<sup>1</sup> | 128 |
| Br4923 | Kyoto-2 | 231 |
| TN | MRHRU-235-G<sup>1</sup> | 110 |
| TN | Kyoto-2 | 249 |
| MRHRU-235-G<sup>1</sup> | Kyoto-2 | 212 |

<sup>1</sup> Strain was later removed from analysis.

### 3.3. Read mapping and SNP calling using 300bp reads

When running Snippy on 300bp reads, over 80% of SNPs were consistently identified correctly as shown in figure 3. Only two FPs were called and occurred in the analyses involving Br4923 as reference. MRHRU-235-G was removed from analysis at this stage due to an issue when calling SNPs; including this strain led to all Snippy output files being empty.

**Figure 3.**
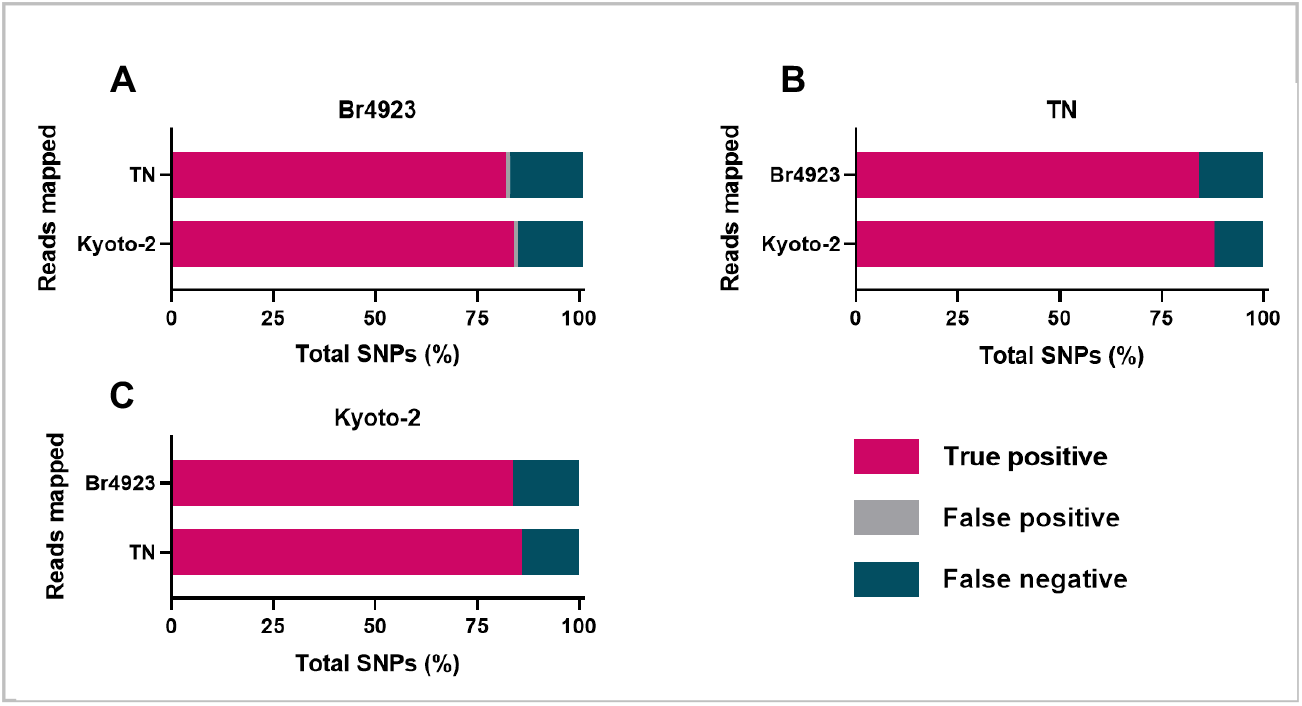
The proportion of ground truth SNPs called as TP (pink), FP (grey) and FN (blue) by Snippy. The proportions are shown per mapped genome (y axis) against (A) Br493, (B) TN, and (C) Kyoto-2 as the reference.

### 3.4. Assessment of clinical data

Of the total 1,112 clinical samples available in the ENA database, across paired and single end Illumina datasets and a variety of other platforms (Supplementary table 1). Only 411 (37%) were retained after mapping filtering. The collated MultiQC report on these samples revealed the most common read lengths as 75bp and 150bp (Figure 4). This information informed the settings used to simulate realistic read lengths with InSilicoSeq.

**Figure 4.**
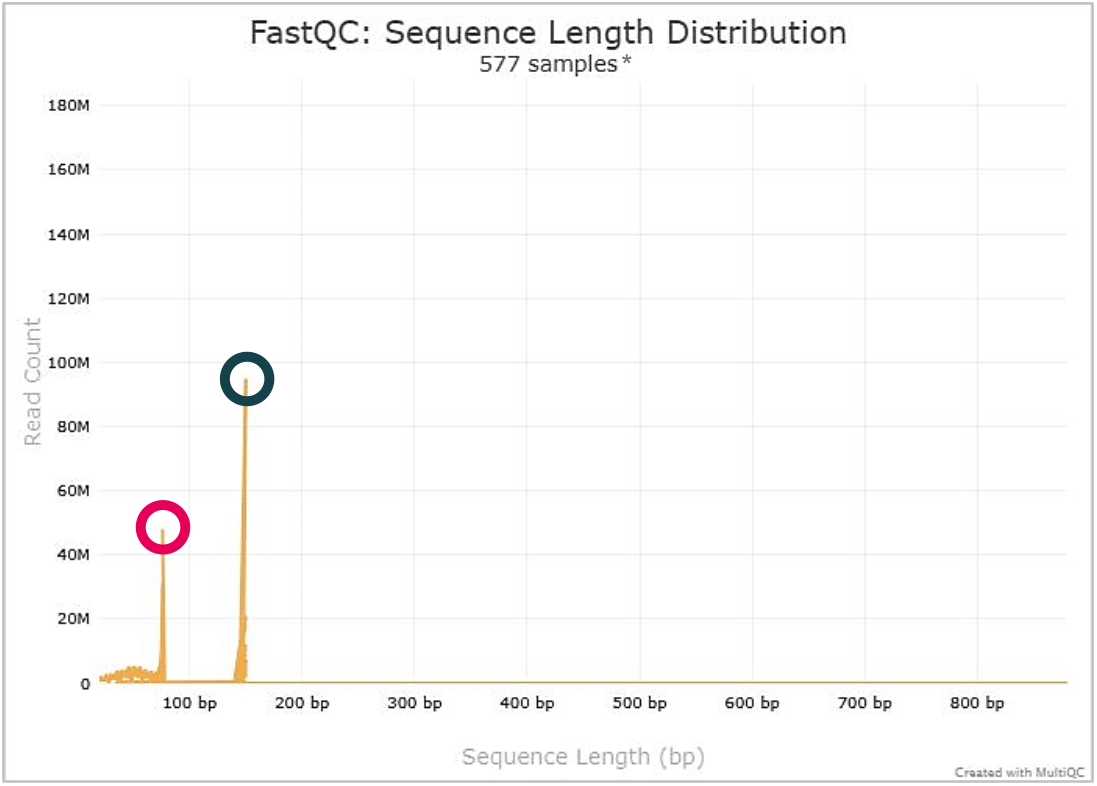
Distribution of short read sequence lengths observed in the clinical data. The most common read lengths were 75bp (shorter peak, circled in pink) and 150bp (taller peak, circled in blue). **\***Paired-end reads are processed separately.

### 3.5. Read mapping and SNP calling using 75 and 150bp reads

With both of these shorter read lengths, Snippy correctly called at least 78% of SNPs (Figure 5); the results are almost identical between the comparisons using both 75bp and 150bp reads. A single false positive was observed in every comparison involving Br4923 as either the reference (Figure 5A) or mapped sample (Figure 5B&C).

**Figure 5.**
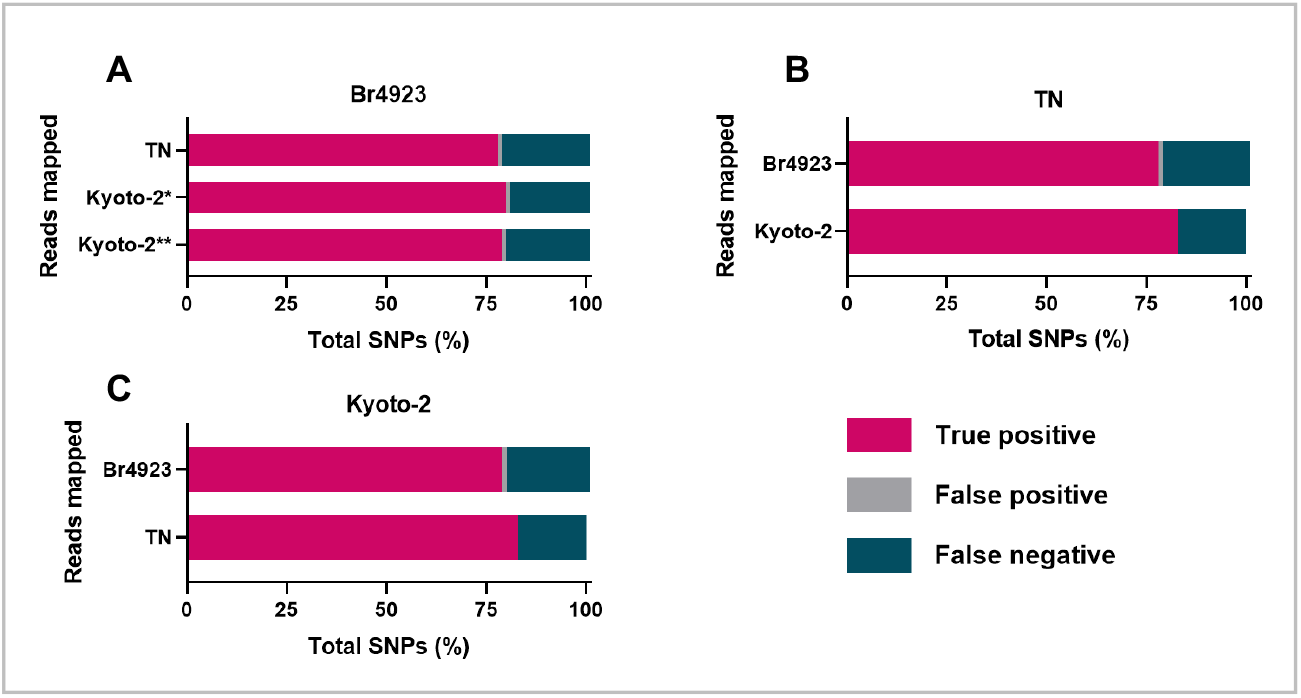
The proportion of ground truth SNPs called as TP (pink), FP (grey) and FN (blue) by Snippy. The proportions are shown per mapped genome (y axis) against (A) Br493, (B) TN, and (C) Kyoto-2 as the reference. The 75 and 150bp reads produced the same results, except where the simulated reads of strain Kyoto-2 were mapped to Br4923. *75bp read results; **150bp read results.

### 3.6. Pipeline performance estimation

Supplementary tables S1 and S2 contain the full confusion matrices of TPs, FPs and FNs for each read length. These tables were used to calculate the precision, recall and F1 score metrics shown in figure 6. Perfect precision was observed for all 300-bp mapping except those involving Br4923 as the reference (Figure 6A). Recall scores for each of the genomes all exceed 0.8, with the F1 score consistently around 0.9 for each comparison. Precision, recall and F1 scores were identical between both 75-and 150-bp lengths except for the recall score in the comparison mapping the Kyoto-2 150bp reads to Br4923 (figure 6D); 0.8 for the 75bp reads and 0.79 for the 150bp reads. The precision scores were lower than 1 in all but two comparisons due to the presence of FPs from Br4923 comparisons.

**Figure 6.**
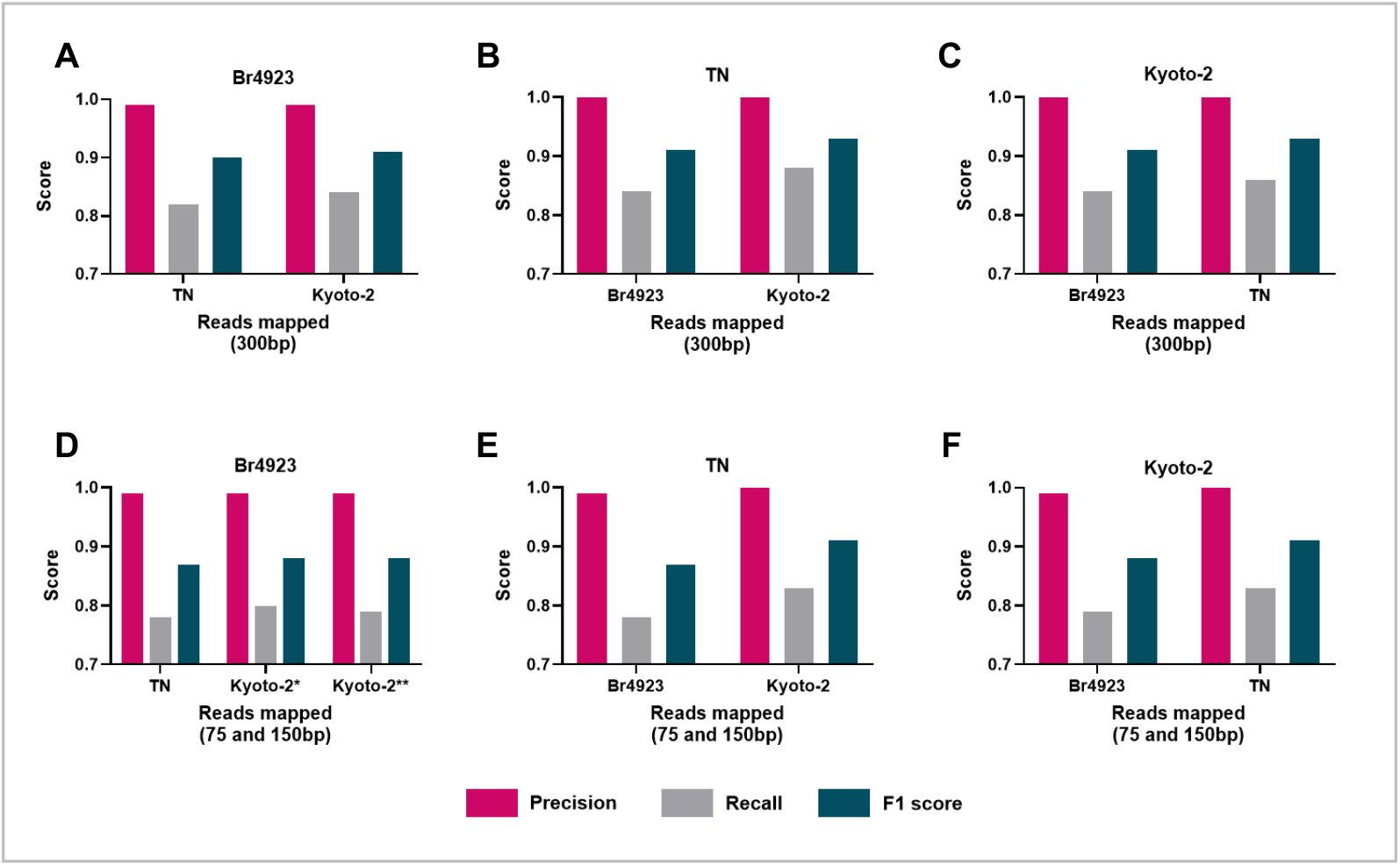
The metrics calculated using the TP, FP and FN SNPs for (A, B, C) 300bp read mapping and (D, E, F) 75 and 150bp read mapping. (A and D) display the results using Br493 as reference, (B and E) show the results of the TN strain as reference, and (C and F) correspond to the analyses using Kyoto-2 as reference. On the y-axis are the simulated short reads for each genome compared against the reference. The 75 and 150bp reads produced the same results, except where the Kyoto-2 simulated reads were mapped to Br4923. *75bp read results; **150bp read results.

### 3.7. Low complexity and repeat region masking

An average of 220 regions (range 219-222) in each genome contained repeat regions (using the K50E4 approach), while around 553 regions were deemed low complexity (Table 2). Using either mask, 3.3-3.4% of the genome was consistently removed. The full number of SNPs masked and the subsequent impact on metrics can be seen in supplementary tables S3, S4, and S5.

**Table 2.** The number of sites and proportion of each genome masked using the k-mer-based repeat mask and low complexity region masking.

|  | Br4923 | TN | Kyoto-2 |
| --- | --- | --- | --- |
| Genome size (bp) | 3268071 | 3268203 | 3268122 |
| **K-mer-based repeat masking** |  |  |  |
| Number of masked regions | 220 | 222 | 219 |
| Total length masked (bp) | 111966 | 112200 | 111958 |
| Genome masked (%) | 3.4 | 3.4 | 3.4 |
| **Low complexity region masking** |  |  |  |
| Number of masked regions | 548 | 559 | 551 |
| Total length masked (bp) | 109276 | 109491 | 109365 |
| Genome masked (%) | 3.3 | 3.4 | 3.3 |

### 3.8 Reference-free distance estimation

Figure 7 shows the ratio of SNPs detected using SKA2 distance estimation on the complete genomes, InSilicoSeq reads and Skesa assemblies compared the ground truth. Between 76% and 81% of SNPs are called through SKA2. These results are consistent across whole genomes, simulated reads, assemblies and read lengths.

**Figure 7.**
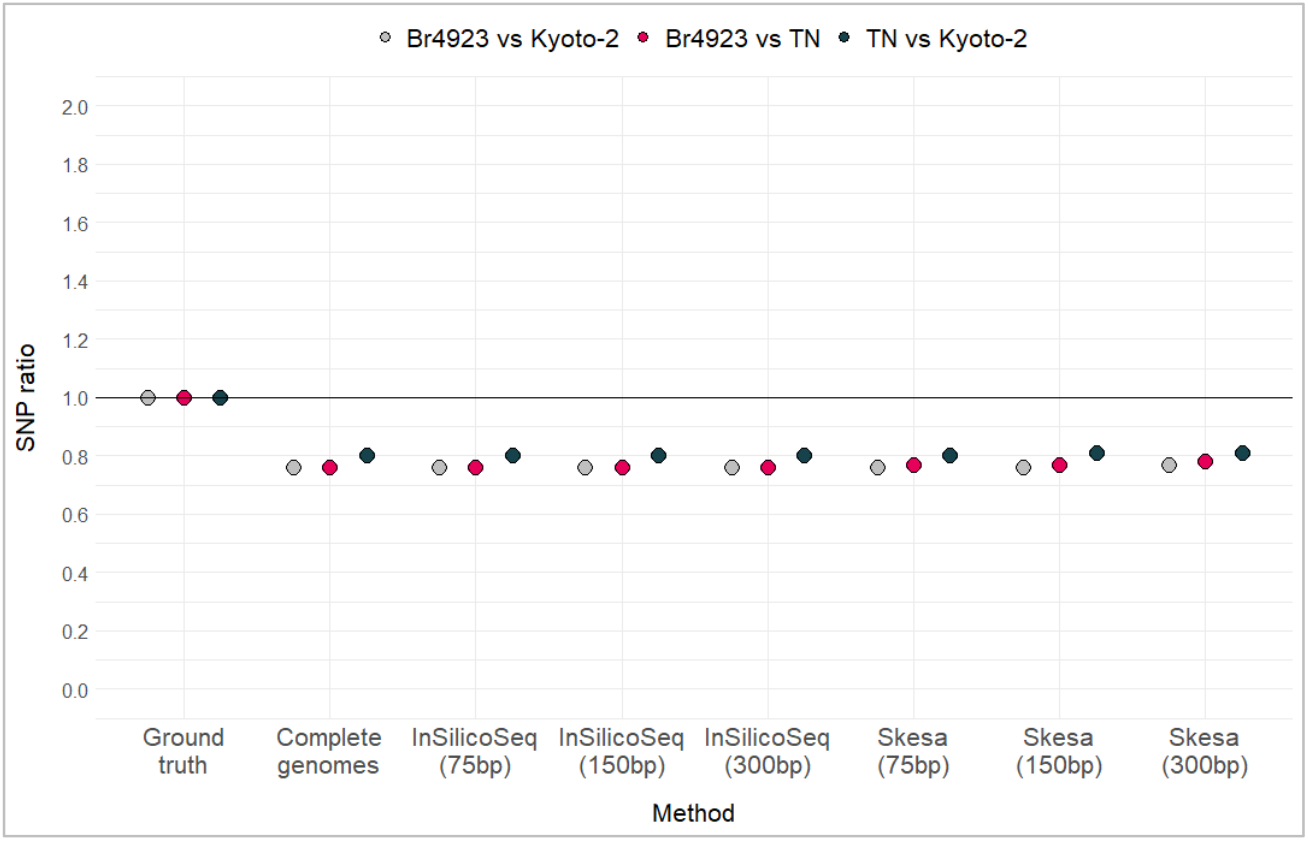
The ratio of SNPs identified using SKA2 distance estimation compared to the ground truth, which is set to 1. SKA2 was performed using the original complete genomes, all simulated reads and Skesa assembled genomes. The read lengths used are shown in parentheses. The colours of each dot correspond to each genome comparison as shown at the top.

## 4. Discussion

### 4.1. Ground truth SNPs

Previous works have already established the extreme clonality and lack of complexity in the M. leprae genome [18,23,24]. The initial findings of this study corroborate this, with less than 250 SNPs making up the ground truth between genomes of 3.3Mb (table 1). For context, M. tuberculosis (Mtb) is considered a low-diversity organism itself but has a SNP frequency rate of 1 in 3 Kb compared to the 1 per 28 Kb of M. leprae [18]. The limited number of SNPs observed between M. leprae strains confirms that the high resolution afforded by SNP calling is needed to achieve differentiation.

The initial ground truth results show that the total number of SNPs between strains does not correlate as clearly to geographical region as SNP typing. The lowest SNP distance observed in this study, 110, predictably occurred in the alignment of two strains of Indian origin – TN and MRHRU-235-G. While the SNP type of MRHRU-235-G is unknown, its geographical proximity to TN (type 1A) suggests it is likely to also be type 1 [23]. Comparison of TN and MRHRU-235-G against the Brazilian strain Br4923 identified 155 and 128 SNPs, respectively. Br4923 is type 4P, so a larger SNP distance would be expected due to the larger genetic distance between type 1 and type 4 strains [23]. Consequently, the results involving Kyoto-2 were surprising. As SNP type 3K, Kyoto-2 is genetically closer to Br4923 [24,58]. However, comparing TN, MRHRU-235-G and Br4923 to Kyoto-2 produced the highest SNP distances of 249, 212 and 231, respectively.

### 4.2. Read mapping

In molecular epidemiology studies, the first genome completed typically becomes the reference genome for subsequent analyses [16-18]. However, as DNA sequencing technologies advance, genomes sequenced using Next Generation Sequencing platforms may be of a higher quality than their first-generation-sequenced counterparts. As the first strains to be completed, TN and Br4923 were sequenced using Sanger sequencing [18,23,29]. MRHRU-235-G and Kyoto-2 were later sequenced using Illumina technology [29]. The downstream effect of using one strain as reference over the others was explored in this study.

To determine whether the reference choice impacts SNP calling, each strain was tested against the others. The original four strains used to establish the ground truth were shredded into short reads to simulate the state of clinical data. When performing SNP calling using these reads, the inclusion of MRHRU-235-G led to the Snippy output files being generated with no data. Removing this strain and re-running Snippy still produced empty files, indicating this could be an issue with the read simulation instead. InSilicoSeq was originally automated to run each read simulation together, so this was re-attempted with MRHRU-235-G reads simulated separately. Unfortunately, the issue persisted. A recent publication has reported that the size of this genome is around 80Kb smaller than other strains, suggesting the coverage or assembly of the strain is incomplete [28]. While this could account for its own lack of results, this does not explain why the Snippy outputs for the other strains were also empty. Ultimately, the strain was removed from subsequent analysis so that the rest of the project could continue within the available time frame.

Upon successfully running Snippy on the three remaining strains, an average 85% of SNPs were correctly called when mapping 300bp reads (table S1).These results did not differ significantly, except for the erroneous calling of two FP SNPs at position 384520 when using Br4923 as reference. The proximity of a SNP to gaps or other SNPs can introduce ambiguity in variant calling. Investigation into these FPs revealed no gaps or other SNPs nearby. Errors may also occur due to read mapping ambiguity introduced by repeat sequences or low complexity regions. However, masking these regions did not remove the FPs. To explore these FPs in more depth, the Mauve GUI was used to visualise the genome alignment between Br4923, TN and Kyoto-2 (supplementary figure 1). This revealed an ambiguous base at position 384520 in Br4923. This was aligned with cytosine at positions 384503 and 384458 in TN and Kyoto-2, respectively. At position 384520 in the other two genomes, all three strains possessed the same base. As a result, SNP calling using the Br4923 produced a FP SNP at this position, whereas the other two strains did not. This highlights the importance of having high-quality sequences for SNP calling in such low diversity genomes.

When mapping reads as short as 75bp, an average 80% of SNPs were still correctly identified (table S2). All results were the same for 75bp and 150bp reads, except for one SNP which was missed between Kyoto-2 and Br4923 using 150bp reads (fig. 5). Though this difference is minute, it could suggest that the shorter 75bp reads are mapping to more regions, slightly increasing the detection of SNPs in these areas. On top of the original FPs at position 384520, two more FPs were introduced with 75bp and 150bp read lengths. These occurred at the same position each time, with one FP at position 1587643 between TN and Br4923, and one at position 1587582 between Kyoto-2 and Br4923. These SNPs were removed when applying either mask, indicating that these positions are located in low complexity sequences which repeat throughout the genome. This would be supported by their absence in the 300bp analysis, where the additional contextual data in longer read lengths reduces the ambiguity in mapping to these regions.

As well as investigating FPs, genome masking was used to identify where SNPs were missed. On average, 15-20% of SNPs are missed by read mapping. Applying the k-mer-based repeat mask removed an average 95% of FNs across all read lengths, while the low complexity mask removed an average of 92%. This indicates that a considerable number of missed SNPs are occurring in regions prone to ambiguity. The remaining FN SNPs after masking showed poor alignment scores in Mauve (example in fig. S2). The lack of coverage in these regions leads to difficulty in mapping reads to these sites, resulting in SNPs being missed. Across the 300bp reads, both low complexity and k-mer-based repeat masking removed 8% of TP SNPs. With the 75bp and 150bp reads, this number drops to 2%. The trade-off in masking FPs but losing TPs in the process is an area explored in the metrics.

### 4.3. Genome completeness

BUSCO was performed retrospectively to determine if discrepancies in the presence of universally conserved genes could account for the misalignments and issues with MRHRU-235-G. It was hypothesised that the significant pseudogenisation of the M. leprae genome could result in missing or fragmented genes, leading to alignment issues and FPs. Additionally, it was considered that orthologs may be present but missed by the program as they do not “look” the same as those present in other Actinomycetes [17]. However, the results generated showed a consistent 92% match across all four strains (fig. 2). Furthermore, N50 scores for each strain covered the entire length of the genome, indicating high contiguity [29]. Ultimately, this suggests that the original FPs which occur and remain after masking could be a natural limitation of read mapping using low complexity genomes. It is possible that resequencing low coverage regions of these genomes could resolve the FPs and the error caused by MRHRU-235-G, though this would be outside the scope of this investigation.

### 4.4. Pipeline evaluation

Precision, recall and F1 score metrics are used to put the TP, FP and FN calls into context [31]. When using 300bp reads, all precision scores are 1 except for the comparisons involving Br4923 as reference, which are 0.99 due to the FPs. With the 75bp and 150bp reads, the comparisons between TN and Kyoto-2 return precision scores of 1 due to the absence of FPs. The other comparisons, each with one FP, have precision scores of 0.99. At no point in this study does precision drop below 0.99, indicating that over 99% of SNPs called by Snippy are true, regardless of the read length used for mapping.

The high precision scores are a strong start, but recall is required to provide added context. This is where differences emerge between the read lengths used. Recall when using 300bp reads is consistently above 0.8, averaging at 0.85. Using 75bp and 150bp reads results in a minimum recall of 0.78 in comparisons involving Br4923 and a maximum of 0.83 in comparisons between TN and Kyoto-2. This means that a minimum 78% of SNPs are recalled using shorter reads lengths, and this number increases to around 85% when using 300bp read lengths.

While the differences in recall between 300, 75 and 150bp reads was statistically insignificant, the decrease at shorter read lengths could translate to a loss of resolution where samples are mistakenly excluded from transmission clusters as they do not reach SNP cut-off thresholds. In the wider context of leprosy, this could result in cases being missed. With the available data, a read length of 300bp produces optimal results and would be recommended for the sequencing of future clinical samples. Longer short read lengths were not tested, in part due to time constraints but also because longer sequencing runs can introduce errors [59], reducing later sequence quality and subsequent pipeline results [43]. Nonetheless, this could be a source of investigation for future studies.

A key element of epidemiological studies is the ability to balance sensitivity and specificity, identifying as many true cases as possible while minimising misdiagnoses [60]. The same is true of SNP calling. So far, metrics indicate that an average 80% of the total SNPs between strains are called, and 99% of these SNPs can confidently be considered true. These figures can be summarised by the F1 score, which balances precision and recall. A score of 1 would indicate 100% recall and perfect precision which is unrealistic for real-world data. Using 300bp reads, the F1 score is consistently 0.9 or above. With the 75 and 150bp reads, F1 ranges between 0.87 and 0.91. Considering the low complexity of *M. leprae* genomes and the read mapping ambiguity this can produce, these results confirm that Snippy is robust, reliably and consistently calling a substantial proportion of SNPs across read lengths of 75 to 300bp.

Masking the genomes does lead to an overall increase in precision, recall and F1 scores, as well as the removal of two FP SNPs (tables S3, S4 and S5). At first glance, these findings might indicate that masking is the solution. However, these metrics are increasing because FNs are being masked, not because more TPs are being identified. On the contrary, a small proportion of TPs are also being masked. While this is not reflected in the metrics, which suggest improved recall, this actually means that more SNPs are being missed. Once again, this decrease in resolution could lead to cases being missed due to exclusion from transmission clusters further downstream. Ultimately, the trade-off discussed earlier between masking FPs but missing TPs is not worthwhile. As such, masking is best to use as an exploratory tool to identify where FPs occur and TPs are missed.

### 4.5. Reference-free distance estimation

The reliance of a reference in both genome alignment and read mapping approaches could introduce bias which can ultimately impact SNP calling [61]. ‘Soft’ reference bias is the term used to describe the increased likelihood of the reference variant being called when the input sequences have poor coverage [62]. ‘Hard’ reference bias occurs when SNPs exist in regions of the input sequences which are absent from or do not match the reference [63]. This could explain the 20% of SNPs missed in read mapping.

SKA2 offers a reference-free alternative, instead splitting sequences into k-mers which flank a central base. Where matching k-mers are present in other sequences, the central bases are compared to determine if variation is present [49]. SKA2 was performed on the original complete genomes and all simulated short reads. An assembly step using Skesa was also performed to investigate whether short read assembly would have any impact on the results. No difference was observed in the number of SNPs detected between the complete genomes or short reads (table S6). Between Br4923 and TN, 76% of SNPs were detected. Likewise, 76% of SNPs between Br4923 and Kyoto-2 were identified. Between TN and Kyoto-2, 80% of SNPs were called. Surprisingly, SKA2 performed slightly better with the assemblies than with reads or complete genomes. This increase could consist of FPs caused by misalignments during assembly. These are typically resolved when preparing a genome for announcement [64], hence the absence of these SNPs in the original genomes and short reads. Ultimately, the addition of an assembly step does not seem to provide any significant increase in resolution.

A limitation associated with using SKA2 is that it does not provide SNP positions. Thus, TP, FP, FNs and metrics cannot be investigated with these outputs. Additionally, it means that core SNPs cannot be distinguished from accessory SNPs for clustering. SKA2 does offer a mapping function which provides SNP positions. However, this requires a reference and does not perform as well as Snippy (table S7). Another known limitation with SKA2 is its inability to recognise SNPs which occur in close proximity [31]. This does not seem to be an issue with the strains tested in this study, as the proportion of SNPs called was comparable to read mapping. Nonetheless, this is worth considering if using SKA2 in future investigations.

### 4.6. Clinical sample clustering

To determine whether the 20% of missed SNPs impacts resolution, this study explored how transmission clustering would look with the available clinical samples. A core SNP alignment file was generated by performing the Snippy ‘multi’ function on the clinical dataset against the TN strain as reference. No established SNP cut-off thresholds exist for *M. leprae* so the established 5 and 12 SNP cut-off distances for Mtb were used as a frame of reference [16]. When analysing these results, it was discovered that all 411 samples were retained across different clusters with both cut-offs. This suggests that the core SNPs within *M. leprae* genomes are reliable and robust, and that the missing 20% may consist of accessory SNPs which do not have a significant impact on sample clustering.

There was an initial attempt to explore whether the core SNP alignment could be used to construct a phylogenetic tree, once again using the TN strain as reference. The intention was to determine whether a phylogeographical correlation could be seen using core SNP distances, but both RAxML-NG and IQ-TREE [65,66] encountered memory limitations. Instead, an alternative approach was taken, mapping all clinical reads to each strain as reference. Clinical samples with no location metadata were removed, and the remaining samples were visualised in scatter graphs using one reference on each axis (figure S3). The intention was to see whether samples from the same location clustered. The plots revealed no particular correlation between SNP distance from the references and geographical origin. While this is consistent with previous reports of SNPs lacking the variability for fine-scale investigations [6,7,18,67], this approach utilised all of the SNPs present between each sample and the reference. This does not reflect the number of SNPs occurring between each sample. Thus, future attempts to visualise this data should concentrate on using the output of the core SNP alignment.

These results conclude that further investigation into transmission clustering is crucial. For instance, *M. leprae* has a much slower SNP frequency rate than Mtb [18]. This could mean that the cut-off thresholds used for Mtb are not stringent enough for *M. leprae*. Secondly, only the TN strain was used as reference in the cut-off-based clustering, posing the question of whether using the other strains would impact the results. Finally, geographical location was not incorporated into the interpretation of the constructed clusters. This was in part due to time constraints, but also because 109 of the 411 clinical samples did not contain information on sample location. This highlights the limitations posed by the current state of *M. leprae* sequence data curation.

### 4.7. Wider implications

This study set out to determine whether a SNP calling pipeline could be used to reconstruct transmission clusters considering the remarkably low complexity of *M. leprae*. Findings indicate that this is indeed possible - both the reference-based and reference-free methods consistently called an average 80% of the total SNPs between strains, and ground truth validation provides a high degree of confidence in these results. Additionally, the pipeline is low-cost. All of the tools used are free and easily available through Conda with thorough documentation. Additionally, all tools are easy to use and simple to automate. Ultimately, this study highlights three key considerations for *M. leprae* SNP calling pipelines.

#### 1. Read mapping provides additional data

While read mapping and distance estimation produce comparable results, the k-mer-based approach of SKA2 does not provide SNP positions. This is a crucial element for clinical molecular epidemiology. Researchers are identifying increasing numbers of high-confidence SNPs which result in drug resistance or differences in disease manifestation [68-70]. Comparing SNP positions to these known loci can allow the accurate prediction of these traits, ultimately influencing the treatment of patients [28]. Furthermore, the ability to examine SNP distances facilitates the reconstruction of transmission clusters as shown with Mtb [71]. This could be implemented with *M. leprae* genomes to enhance surveillance efforts. SKA2 may therefore be better suited to rapid preliminary investigations [49]. Alternatively, SKA2 could be implemented on isolates within SNP types or subtypes. An investigation into critical pathogens using SKA2 found that the estimation of SNP distances could provide added resolution to identify outbreaks in localised regions [72]. This may not be applicable to *M. leprae* due to its low diversity but would certainly be worth further examination.

#### 2. Reference genome selection is important

A major limitation in *M. leprae* studies is the lack of available complete genomes. The combination of its slow growth, uncultivability and lack of research funding as a neglected tropical disease means that researchers typically focus on amplifying targeted sequences of interest instead of assembling and annotating whole genomes. This means that the conclusions drawn from the results of this study are limited and may evolve with the incorporation of more available genomes in the future. Many questions arise from this implication. For instance, would Br4923 consistently call FPs across a wider set of data? Could the inclusion of more complete genomes lead to FPs called using the TN and Kyoto-2 strains? Would phylogeographic patterns begin to emerge? Nonetheless, all three genomes performed comparably and introduced minimal errors, making them suitable as references for read mapping approaches. Additionally, the similarity in SNP calling using SKA2 indicates that reference bias may not be a significant factor in these analyses. Another consideration is the reliance on one reference genome for the basis of studies across different SNP types. It has been documented that the H37Rv reference genome of Mtb is a lineage 4 strain, and may not be optimal to use for other, more genetically distance lineages [16]. Thus, it would be worth exploring whether the use of a type-specific reference increases SNP recall, accuracy and overall resolution.

#### 3. High quality data is crucial

Perhaps the most important takeaway from this study is the reliance of pipelines on high quality data. To begin with, the original issue with strain MRHRU-235-G led to its removal from analysis, taking the total number of useable genomes down to three. The strain is hypothesised to be incomplete due to its reduced size compared to other strains [28]. As the foundation of this pipeline, complete genomes need to be high quality in order to produce reliable results downstream. Secondly, a key point to highlight is the filtration of 1,112 supposed *M. leprae* sequences down to 411. The removed samples had poor coverage, contained host DNA contamination or were simply mislabelled primer sequences. Despite this, FastQC and MultiQC performed on the remaining 411 samples revealed high Phred scores of around 35. Of these samples, 109 did not provide location data. Metadata, particularly the sample location, provides crucial context for molecular epidemiology studies and its absence limits the interpretation of results. Ultimately, thorough contextual data and accurate results are needed in order for a molecular epidemiology pipeline to be effective.

### 4.8. Future directions

Among the recommended investigations already discussed, there a number of other explorations which could solidify this SNP calling framework as a standard approach for clinical molecular epidemiology. Firstly, investigating whether this pipeline provides enough resolution to distinguish mixed infections and determine whether persisting cases are relapses or reinfections [73-75]. If sufficient, this could provide insights into local transmission and how best to treat patients with complex infections. Additionally, it would be useful to examine how portable this pipeline is to other species. In particular, *M. lepromatosis* – a secondary (but less prolific) cause of leprosy discovered in 2008 [76,77] An adaptable pipeline would further enhance leprosy surveillance by identifying cases caused by both species. Finally, as many studies have suggested, it may be worth investigating the combination of SNP calling stability and VNTR diversity to maximise resolution [3,78,79].

## Conclusions

*M. leprae* is one of the most uniquely challenging pathogens to study, owing to its diminished genome, low complexity, and inability to grow on culture media. Advances in the ability to extract and sequence DNA from skin lesions and biopsies has facilitated major breakthroughs, such as the development of a robust SNP-based strain typing system for phylogeographical investigations. The eradication of leprosy relies on detecting transmission clusters to inform targeted control strategies. Declines in case detection and reports of emerging drug resistance highlight the growing need for a clinical molecular epidemiology pipeline.

This study demonstrates that both read mapping and distance estimation approaches can achieve accurate SNP calling. Thus, a clinical pipeline for *M. leprae* transmission clustering is feasible. Future studies should build upon these findings by combining epidemiological data with SNP distances to identify cut-off thresholds for clustering. Additional directions include investigating the efficacy of SNP calling with mixed infections, relapses and reinfections to guide clinical decision-making and identify local transmission events.

## Acknowledgements

The author would like to thank supervisors Dr Conor Meehan and Dr Maria Rosa Domingo Sananes as well as collaborators Dr Daniel Whiley and Aureliana Chilengue for their support and guidance during this study.

## Data Availability Statement

The original complete genomes used can be found at the National Center for Biotechnology Information Genome database (GenBank accession numbers GCA_003253775.1, GCA_000195855.1, GCA_003584725.1, GCA_000026685.1). All *M. leprae* clinical samples can be obtained from the European Nucleotide Archive. Supplementary data tables, pipeline scripts, clustering output files and the clinical sample MultiQC report can be accessed <u>**here**</u>.

