## Supplementary table for "Evaluating a SNP calling pipeline for *Mycobacterium leprae*"

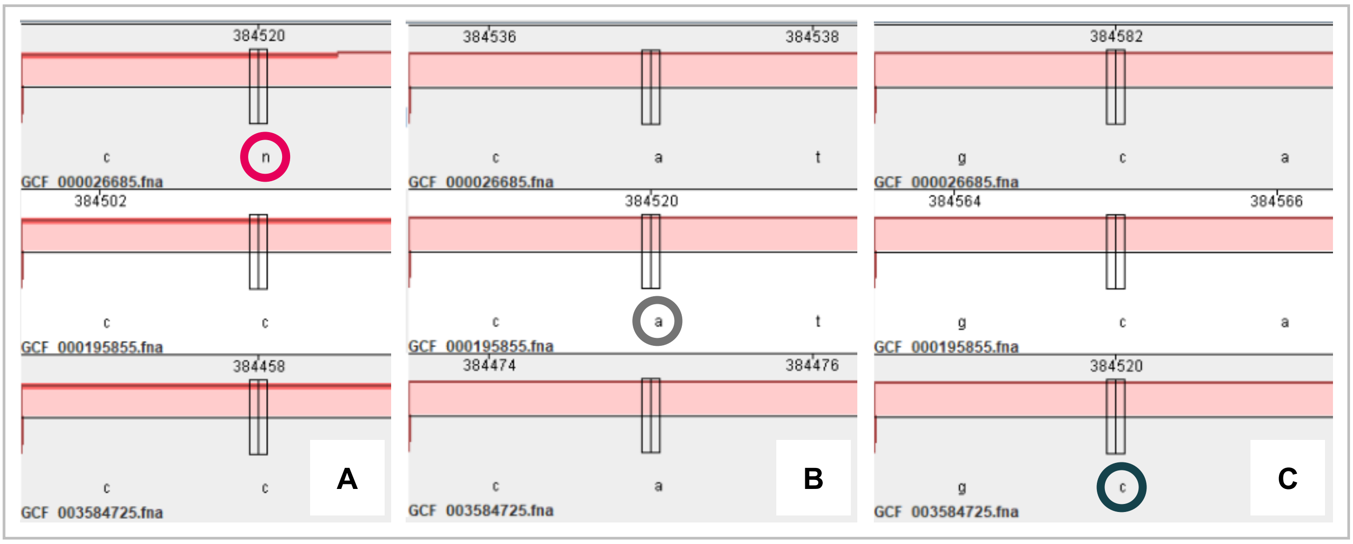


**Figure S1.** Alignment between (A, top row) Br4923, (B, middle row) TN and (C, bottom row) Kyoto-2. At position 384520 in Br4923, an ambiguous base occurs (circled pink), while the other two strains show cytosine. At position 384520 in TN, all bases are adenine, while cytosine appears in all bases at position 384520 in Kyoto-2.

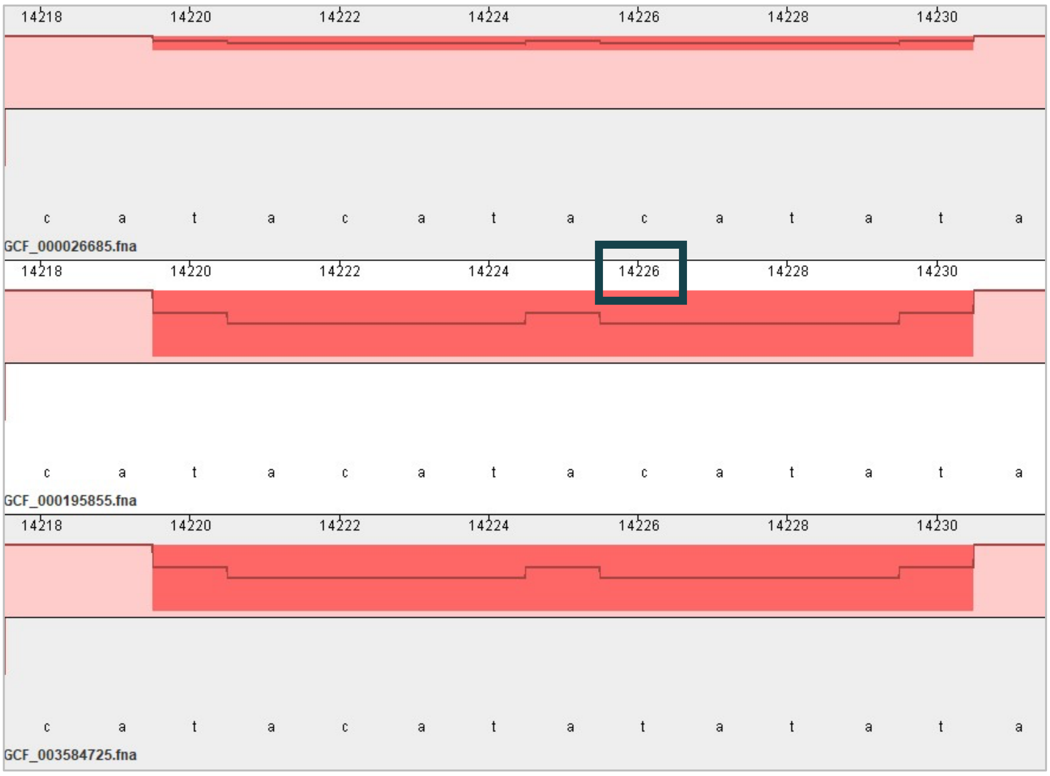


**Figure S2.** Alignment between Br4923 (top row), TN (middle row) and Kyoto-2 (bottom row). A SNP occurs in the Kyoto-2 strain at position 14226 (blue box). The red background and dip in the alignment conservation graph indicates a region of poor alignment and low coverage.

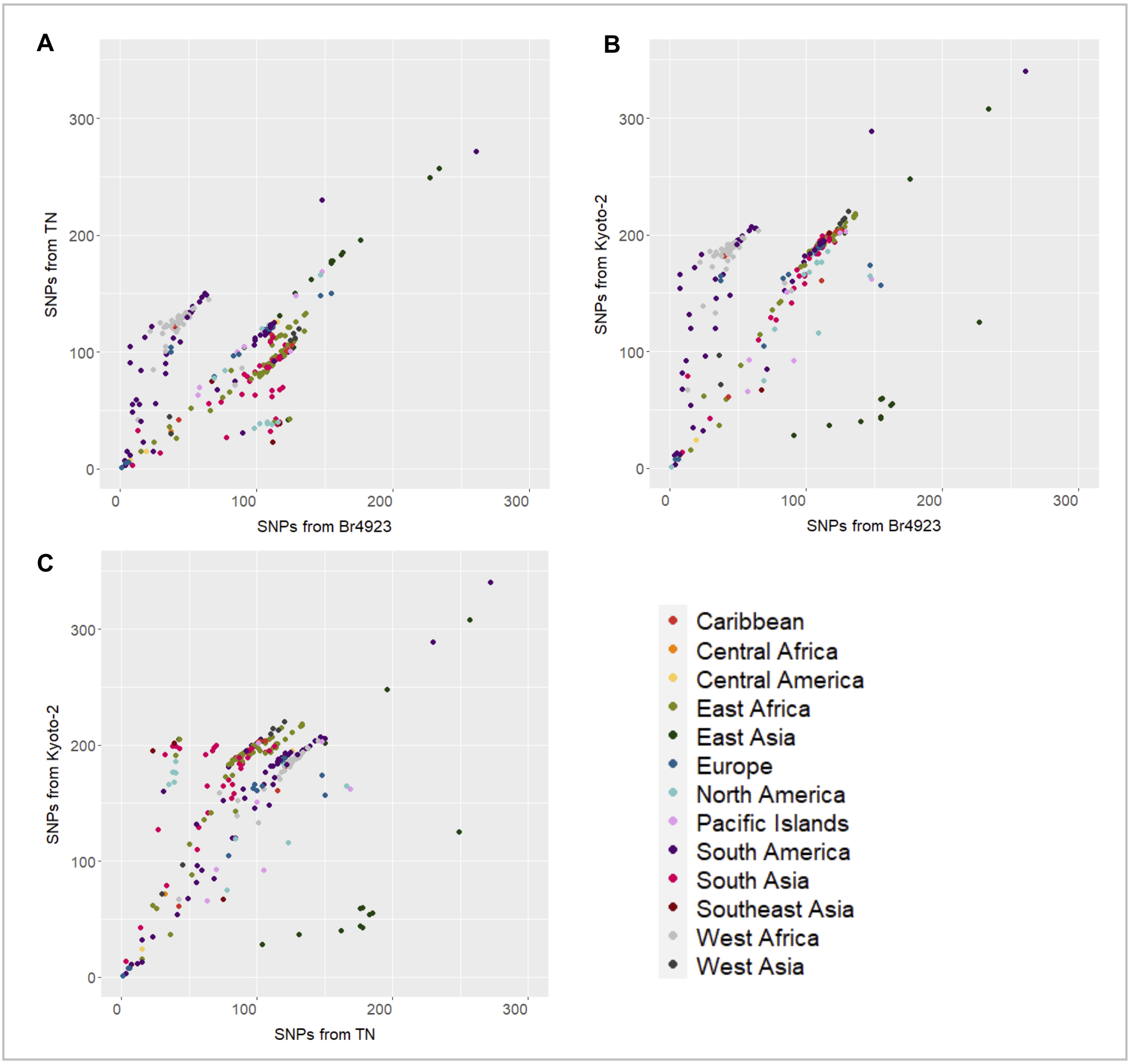


**Figure S3.** SNPs between clinical samples with location metadata and their SNP distances from the (A) Br4923 and TN strains, (B) Br4923 and Kyoto-2 strains, and (C) TN and Kyoto-2 strains. Data is coloured based on geographical location for easier visualisation of clustering.
